# Joint reconstruction of multi-contrast relaxation-diffusion imaging

**DOI:** 10.64898/2026.10.21.760608

**Authors:** Yunqi Wang, Qiang Liu, William Consagra, Lipeng Ning, Kawin Setsompop, Yogesh Rathi

## Abstract

Diffusion MRI across multiple gradient directions, b-values and echo times (TEs) provides complementary tissue contrast but requires extensive sampling. Joint k-space and q-space undersampling requires reconstruction that preserves diffusion- and TE-dependent signal variation while exploiting shared anatomical information. Here we introduce DIFFUSE (Diffusion-relaxometry Implicit Field Fitting from Undersampled *k*–*q* Space across Echoes), a subject-specific joint *k*–*q*–TE image reconstruction framework. It fits a single implicit neural representation (INR) directly to complementary undersampled multi-coil *k*-space measurements across diffusion encodings and echo times. The INR (a self-supervised neural network) maps spatial location, b-value and TE to baseline b0 signals and spherical harmonic coefficients for directional diffusion-weighted image reconstruction. The measurement-specific forward model incorporates phase, coil sensitivities and motion and eddy current related transforms. We evaluated DIFFUSE using retrospective Human Connectome Project data, controlled motion simulations and in vivo multi-shell and multi-TE acquisitions. At sixfold in-plane acceleration, DIFFUSE reduced normalized mean squared error by 57–73% relative to SENSE across two diffusion shells in the retrospective experiment. In the controlled motion experiment, motion-aware DIFFUSE reduced normalized mean squared error by 66–72% relative to static DIFFUSE. The prospective study demonstrated multi-shell reconstruction at 1.3 mm isotropic resolution with significantly high signal-to-noise ratio (SNR). In a separate multi-TE experiment, a single representation jointly reconstructed all nine contrasts corresponding to 3 b-value/TE combinations. DIFFUSE thus provides a common reconstruction strategy for accelerated protocols spanning multiple diffusion encodings and echo times.

## 1. Introduction

Diffusion MRI (dMRI) probes tissue organization by measuring signal attenuation under different diffusion encodings. Sampling multiple gradient directions supports tensor and orientation-based descriptions of anisotropic tissue architecture (Basser and Pierpaoli, 1996; Westin et al., 2002; Tuch, 2004; Descoteaux et al., 2007; Tournier et al., 2007). Multiple *b*-value shells provide complementary sensitivity to tissue microstructure (Zhang et al., 2012; Consagra et al., 2025), while variation in echo time (TE) adds relaxation-dependent contrast for joint diffusion–relaxation analysis (Veraart et al., 2018; Ning et al., 2020). A protocol spanning direction, *b*-value, and TE can therefore characterize several aspects of tissue within one examination. Each combination nevertheless produces a distinct diffusion-weighted image (DWI) contrast and requires corresponding multi-coil measurements. The number of contrast-specific acquisitions grows rapidly as these protocol dimensions are combined, lengthening the examination and increasing exposure to subject motion. High-resolution multi-shot dMRI can improve structural-network characterization (Merenstein et al., 2023), and in-plane segmented multi-slab acquisition with denoiser-regularized reconstruction has enabled submillimeter 3D dMRI (Li et al., 2026).

This acquisition burden arises in two sampling domains. Here, *q*-space denotes the diffusion-encoding domain defined by gradient direction and *b*-value, whereas *k*-space denotes the spatial Fourier domain measured by the receiver coils. A practical protocol may retain only a prescribed subset of diffusion encodings (Chen et al., 2021) and may also acquire incomplete *k*-space for each retained encoding/TE condition. Complementary masks distribute Fourier samples across related conditions, so locations omitted for one contrast may be observed for another. The reconstruction target is the set of contrasts specified by this multidimensional protocol. The resulting task is therefore broader than reconstructing one undersampled image: incomplete multi-coil Fourier observations must be integrated across *q*-space and TE without erasing the differences that define each contrast.

Parallel imaging and compressed sensing reduce the number of Fourier samples required for each image through coil encoding, autocalibration, or image sparsity (Pruessmann et al., 1999; Griswold et al., 2002; Lustig et al., 2007). Calibration information can be shared across volumes through a common GRAPPA kernel, while separate image-reconstruction objectives treat each diffusion- and TE-dependent contrast independently. The corresponding DWIs depict the same anatomy, with shared boundaries and spatial organization, while their intensities vary systematically with gradient direction, *b*-value, and TE. Complementary sampling enables a shared signal representation to exploit this common spatial structure across conditions. Conditioning the representation on the acquisition variables preserves the diffusion and relaxation information specific to each contrast.

Cross-measurement reconstruction provides a way to exploit the common structure of multidimensional dMRI without treating every DWI as an isolated inverse problem. Joint *k*–*q* reconstruction and direct ensemble average propagator recovery show that diffusion-domain relationships can constrain incomplete measurements, although these approaches impose different priors and recover different objects (Wu et al., 2019; Ye et al., 2025; Mani et al., 2021; Sun et al., 2019). Multi-TE diffusion studies likewise exploit echo-dependent information through diffusion–relaxation modeling, acquisition design, or joint *k*–TE reconstruction (Ning et al., 2020; Ji et al., 2022a; Liu et al., 2024; Ji et al., 2022b; Jun et al., 2026). These advances address complementary parts of the problem, but they do not by themselves provide a shared signal representation spanning direction, *b*-value, and TE linked directly to acquired multi-coil data. These requirements motivate a joint *k*–*q*–TE reconstruction setting in which diffusion-encoding and TE dimensions are coupled within one reconstruction objective.

A subject-specific continuous field can provide this shared representation over spatial and acquisition coordinates. For joint dMRI, the field must encode the diffusion- and TE-dependent signal itself rather than only a downstream orientation distribution or compartment parameter. It must also be constrained directly by acquired multi-coil *k*-space samples, rather than by images produced in a preceding reconstruction stage. Existing dMRI implicit neural representations (INRs) have mainly been fitted to reconstructed diffusion images (Consagra et al., 2024; Hendriks et al., 2026; Zeng et al., 2026), whereas acquisition-domain INRs in quantitative MRI demonstrate direct *k*-space fitting (Lao et al., 2026). Combining a diffusion-signal field with acquisition-domain consistency yields a single per-examination representation that preserves measurement-specific contrast variation while allowing complementary measurements to constrain shared parameters jointly. Cross-measurement sharing is valid only when acquisition-specific effects remain outside the tissue field and every measurement is related to a common reference-space representation. Shot-dependent phase alters complex *k*-space, while coil sensitivities and sampling masks define how each measurement is encoded. Rigid head motion breaks this correspondence by changing the mapping between acquisition and reference-space coordinates.

DIFFUSE (Diffusion-relaxometry Implicit Field Fitting from Undersampled *k*–*q* Space across Echoes) implements a subject-specific joint reconstruction. It parameterizes a five-dimensional field over spatial position, *b*-value, and TE, and represents directional dependence through real spherical harmonic (SH) coefficients. Directional DWIs are reconstructed by evaluating the SH basis at the corresponding diffusion-encoding direction. Baseline channels represent the direction-independent *b* = 0 signal. The shared signal field generates each contrast, which is then phase modulated, encoded by the corresponding coil sensitivity, Fourier transformed, and restricted to acquired locations by its sampling mask. When direction-specific affine spatial transforms are available from head-motion and eddy-current correction, their coordinate mappings and corresponding *b*-vector reorientations are applied within the measurement operator. The INR parameters are fitted anew to acquired multi-coil *k*-space samples for each examination, requiring neither population-level training nor a prescribed biophysical compartment model.

## 2. Related Work

### 2.1. Cross-measurement reconstruction in diffusion and relaxation dimensions

Joint 6D *k*–*q* compressed sensing recovers diffusion signals and the EAP from simultaneously undersampled *k*-space and *q*-space (Cheng et al., 2015), and XQ-SR exploits image-domain *x*–*q* structure for spatial–angular super-resolution (Chen et al., 2019). Non-parametric Gaussian-process models predict single- and multi-shell diffusion signals directly in *q*-space (Andersson and Sotiropoulos, 2015). DAGER introduced Gaussian-process predictions over *q*-space as a prior for joint *k*–*q* reconstruction (Wu et al., 2019). Its multi-shell extension models angular and radial relationships through cross-shell covariance. It also applies a variance-stabilizing correction when fitting Gaussian processes to Rician-distributed magnitude data (Ye et al., 2025). qModeL follows a different route: a denoising autoencoder trained on biophysically simulated *q*-space dictionaries supplies a plug-and-play signal-manifold prior within phase-compensated model-based reconstruction (Mani et al., 2021). Direct ensemble average propagator (EAP) recovery changes the target representation by mapping undersampled (*k, q*)-space to the EAP without first recovering a complete DWI series (Sun et al., 2019). For its in vivo demonstration, undersampled Fourier measurements were retrospectively formed from image-domain HCP data rather than acquired as raw multi-coil data. Together, these methods establish several ways to exploit angular and shell structure, but they use strong priors, whether deep learning based or specific model-based (EAP).

Multi-TE dMRI has developed through related but distinct efforts in signal modeling, acquisition design, and reconstruction. Joint relaxation–diffusion moments use signal variation across diffusion encoding and TE to probe neurite microstructure (Ning et al., 2020). Relaxation–diffusion spectrum imaging provides a complementary joint diffusion–*T*_2_ representation for probing tissue microarchitecture (Wu et al., 2023). Time-division multiplexed simultaneous multi-slice and related TDM sequences reduce the acquisition cost of collecting multiple *b*-value/TE combinations (Ji et al., 2022a; Liu et al., 2024). Joint *k*–TE reconstruction exploits complementary sampling through joint-GRAPPA interpolation across echoes or joint structured low-rank regularization (Ji et al., 2022b). PRIME (Jun et al., 2026) is primarily a phase-reversed interleaved multi-echo acquisition strategy. Its additional echoes support field-map estimation, phase or motion navigation, or diffusion relaxometry according to the acquisition configuration. These studies establish the value of echo-dependent information and cross-echo acquisition structure. Different TEs do not provide repeated observations of the same contrast. Relaxation weighting changes signal amplitude and tissue contrast, so a shared reconstruction must retain TE as a conditioning variable. All these methods use joint reconstruction across the *k*–TE domain in a slice-by-slice manner.

### 2.2. Continuous subject-specific representations and Fourier-domain consistency

INRs replace a discrete collection of independently parameterized voxels or images with a continuous function evaluated over spatial coordinates, acquisition coordinates, or both.

In the subject-specific settings reviewed here, such a field can share information within one examination without population-level training, but its interpretation depends on the output and the observations used for fitting. Neural orientation distribution function fields estimate spatially continuous orientation distributions and their uncertainty from reconstructed signals on a fixed *b*-shell (Consagra et al., 2024). Subject-specific dMRI INRs estimate Standard Model parameters and fiber orientation distributions from reconstructed diffusion images (Hendriks et al., 2026). MINeR uses a direction-modulated INR for image-domain angular interpolation from a sparse set of diffusion directions (Zeng et al., 2026). These methods act on reconstructed diffusion signals and recover either a derived representation, biophysical parameters, or a densely sampled image series.

Direct Fourier-domain fitting provides a different connection between a continuous field and the acquisition process. In multichannel dMRI, reconstruction and denoising choices influence the noise floor, effective spatial resolution, and downstream fiber-orientation estimates (Sotiropoulos et al., 2013; Huynh et al., 2024; Manzano Patron et al., 2024). In multiparametric quantitative MRI, a motion-aware subject-specific INR has been embedded in acquisition-specific Fourier operators and optimized against *k*-space (Lao et al., 2026). This formulation provides a precedent for maintaining acquisition-domain consistency when a subject-specific field is observed at different poses. For joint dMRI reconstruction, however, the represented object must additionally preserve direction- and *b*-dependent signal variation across the acquisition protocol. This comparison separates three design axes for joint dMRI: the represented object, the domain of data fidelity, and the role of acquisition pose in the consistency model.

### 2.3. Measurement-specific phase and motion modeling

Measurement-specific phase can make multi-shot *k*-space inconsistent even when the underlying diffusion magnitude is shared. Phase-aware methods address this inconsistency through explicit phase–magnitude separation, structured low-rank constraints, or learned optimization. MUSE combines coil sensitivity encoding with shot-dependent phase estimates for high-resolution multi-shot DWI reconstruction, and 3D-MB-MUSE extends this formulation to multi-slab, multi-band, multi-shot acquisitions (Chen et al., 2013; Bruce et al., 2017). Self-navigated 3D dMRI combines optimized CAIPI sampling with structured low-rank navigator estimation (Li et al., 2025). PAIR estimates shot phase and a shared magnitude image using a *k*-space low-rank phase prior together with an image-domain weighted total-variation regularizer (Qian et al., 2023). DONATE accelerates structured Hankel recovery with scan-specific adaptive subspaces designed for high shot counts (Qian et al., 2025). PAIR-Net is a supervised model-based iterative network that alternates magnitude and phase updates with multi-coil *k*-space data consistency (Lin et al., 2025). Although their optimization strategies differ, these methods treat shot-to-shot complex modulation as distinct from the shared diffusion-weighted magnitude. Rigid pose and geometric distortion define separate correspondence problems. RPG-MUSE combines reversed-polarity gradients with MUSE to correct *B*_0_- and eddy-current-induced distortions in high-resolution diffusion MRI (Bruce et al., 2018). Pipelines such as FSL eddy estimate head motion together with eddy-current distortions (Andersson and Sotiropoulos, 2016) during post-processing once images have been reconstructed.

### 2.4. Our Contributions

Existing methods either perform joint *k*–*q* reconstruction (but not joint diffusion-relaxation reconstruction), or joint *k*–TE reconstruction (discarding the diffusion directions). Some INR-based methods either work completely in image space (not from the raw *k*-space measurements) or perform reconstruction of relaxometry data alone. Finally, motion correction and eddy-current correction are typically left as post-processing steps in most frameworks. DIFFUSE addresses all these gaps through a unified formulation that fits a subject-specific diffusion- and TE-dependent signal field directly to complementary under-sampled multi-coil *k*-space while accounting for measurement-specific phase and direction-specific affine spatial transforms.

The main contributions of this work are summarized as follows:

1. We formulate image reconstruction as a single subject-specific inverse problem fitted jointly to complementary multi-coil *k*-space across diffusion encodings, TEs, and shots.
2. We introduce a five-dimensional diffusion-signal field whose baseline channels and SH coefficients support continuous directional DWI reconstruction over spatial position, *b*-value, and TE.
3. We construct a phase-aware, measurement-specific forward model that links the shared signal field directly to sampled multi-coil *k*-space incorporating the effects of direction-specific affine spatial transforms for head motion and eddy-current distortion, including rotational *b*-vector reorientation.
4. We evaluate DIFFUSE in retrospective multi-shell acceleration, a controlled rigid-motion stress test, prospective 1.3 mm isotropic multi-shell imaging, and in vivo TDM-EPI spanning three *b*-values and three TEs. The first two settings provide reference-based evidence for validation purposes. The latter two assess high-resolution feasibility and joint representation across acquired conditions without reference images.

## 3. Methods

### 3.1. Problem Formulation

Let *l* ≡ (*z, s, d, b*, TE) denote the combined index for slice, shot, diffusion direction, b-value, and echo time, respectively, while *c* indexes the receiver coils. The k-space data-consistency loss can be written as:

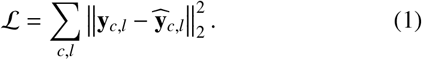

Here, y_*c,l*_ and 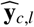 are the measured and predicted k-space data for the same coil and combined index. The latter is generated as:

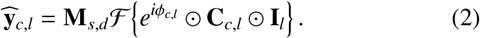

Here, I_*l*_ is the magnitude image, C_*c,l*_ denotes the coil-sensitivity magnitude, and the coil-sensitivity phase is incorporated into 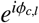. The spatial Fourier transform F maps the coil image to k-space, after which the mask M_*s,d*_ retains only the acquired samples. The symbol ⊙ denotes element-wise multiplication.

The magnitude image is obtained by querying a motion-free reference-space field at the coordinates associated with the acquired slice:

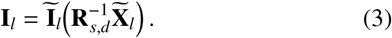

Here, 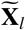 contains all coordinates in the motion-free slice indexed by *l*. The *z* component of *l* selects the slice, while the remaining components identify its acquisition condition. The transform **R**_*s,d*_ denotes a direction-specific affine spatial transform for each shot–direction pair, accounting for head motion and eddy-current distortion. 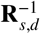 maps motion-free coordinates to the corresponding acquisition coordinates. The rotational component of **R**_*s,d*_ reorients the acquisition-frame diffusion direction into the motion-free reference frame according to 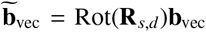. Translation affects only the spatial coordinates. The physical b-value and echo time remain unchanged under the spatial transform. For an acquisition without head motion or eddy-current distortion, **R**_*s,d*_ is the identity and the b-vector is unchanged. The direction-specific affine spatial transforms are estimated by registering the SENSE or initial-INR images used for initialization.

Let 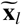 denote a five-dimensional query associated with the motion-free slice indexed by *l*, and let 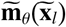, parameterized by the INR weights *θ*, denote the continuous SH coefficient field evaluated at that query. The direction-dependent signal is synthesized as:

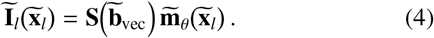

Here, 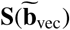 denotes the order-*L* real symmetric SH synthesis row (Descoteaux et al., 2007), and 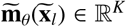 contains the *K* SH coefficients at 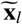. Evaluating Eq. (4) at all spatial locations in the slice yields the motion-free DWI image 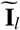 used in Eq. (3). Measurements acquired at *b* = 0 are represented separately by a direction-independent baseline signal field. The INR jointly parameterizes the baseline signal field and the SH coefficient field. Measurements with complementary masks are coupled through the shared continuous field, while each residual is evaluated using its own sampling mask. During network training, the INR parameters *θ* are optimized for subject-specific fitting, whereas MRI experimental parameters such as sampling masks, b-values, gradient directions, and echo times are known.

### 3.2. Implicit neural representation

Figure 1 summarizes the coordinate-to-output branch of the shared five-dimensional INR within the full reconstruction pipeline. The INR takes the five-dimensional coordinate 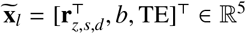 as input. Here, **r**_*z,s,d*_ is the normalized reference-space location associated with slice *z*, shot *s*, and diffusion direction *d*. The *b* and TE entries are the corresponding physical acquisition values used directly. Let *D*_*γ*_ denote the number of Fourier frequency vectors. A Gaussian random Fourier-feature projection 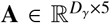 (Tancik et al., 2020) en-codes the input as:

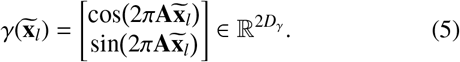

**Fig 1.**
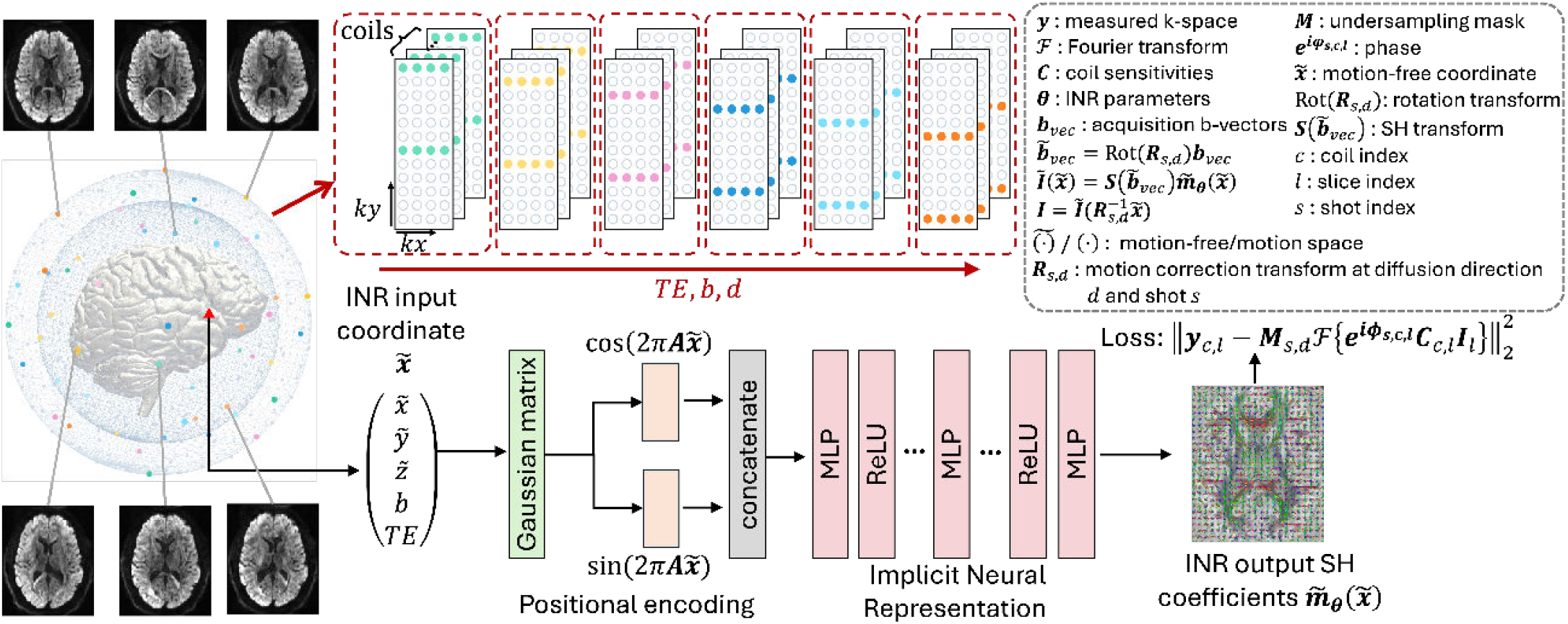
Overview of DIFFUSE for joint k–q–TE reconstruction. Multi-shell, multi-TE diffusion measurements distributed over q-space (left) are acquired with complementary k-space masks across the related measurements defined by each protocol (top). Gaussian random Fourier features encode the five-dimensional INR input comprising three normalized spatial coordinates and physical b-value and TE values before processing by a multilayer perceptron (bottom). The coil index and combined acquisition label select the corresponding phase map, coil sensitivity, sampling mask, direction-specific affine spatial transform, and k-space measurement in the forward model. The INR predicts baseline (*b* = 0) signal channels and SH coefficients of the diffusion signal for directional image synthesis and subsequent diffusion analysis (right). For measurements affected by motion or eddy-current distortion, 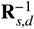 maps motion-free slice coordinates into the corresponding acquisition coordinates, while the rotational component of R_*s,d*_ reorients the b-vectors into the motion-free reference frame before SH synthesis. The reconstructed image is subsequently phase modulated, coil encoded, Fourier transformed, and sampled with the corresponding mask. The displayed loss is the k-space data-consistency term in Eq. (1), with the predicted measurement defined in Eq. (2).

In all experiments, *D*_*γ*_ = 5,000. The rows of A were drawn independently from a zero-mean Gaussian distribution with standard deviation 4.0, and the resulting projection was held fixed. Concatenating its cosine and sine responses produced 10,000 features per query. The shared multilayer perceptron (MLP) *f*_*θ*_ used five fully connected hidden layers, each with 1,024 units and a ReLU activation. A final linear layer produced *n*_0_ + *K* continuous-field outputs, where *n*_0_ denotes the number of direction-independent *b* = 0 baseline signal outputs and *K* denotes the number of real symmetric SH coefficients:

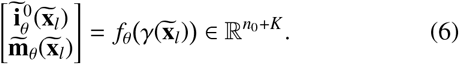

Here, 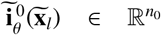 denotes the motion-free, direction-independent baseline signal outputs at *b* = 0. The vector 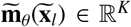 contains the SH coefficients used in Eq. (4). No skip connections or normalization layers were used. With SH order *L* = 8, corresponding to *K* = 45 real symmetric SH coefficients, the MLP contained 14,486,574 trainable parameters for *n*_0_ = 1 and 14,487,599 for *n*_0_ = 2, the two output settings used across the experiments. The fixed 5,000 × 5 Gaussian projection matrix was excluded from these counts. The coordinate encoding and MLP architecture were adapted from the dMRI INR framework of Hendriks et al. (2026).

In this work, our reconstruction framework involved two stages of INR optimization. In the first stage, the INR was initialized with a SENSE reconstructed k-space, termed as “warm-up”, followed by transition to pure undersampled k-space based measurement loss. This strategy was used to reduce the computational burden leading to faster convergence. In the next stage, optimization proceeded using a fresh INR (weights randomly initialized) but with the previous stage INR estimate as the “warm-up” k-space followed by using the actual undersampled measurements for loss calculation. This 2-stage “unrolled” INR optimization framework ensured high data fidelity without getting trapped into the SENSE provided initialization.

## 4. Experiments and Results

We evaluate DIFFUSE in four experiments: 1) performance under various in-plane acceleration factors for undersampled q-space multi-shell data, 2) evaluation of head motion correction, 3) evaluation using real-world prospective 1.3 mm isotropic resolution data with various in-plane acceleration factors, and 4) joint reconstruction of multi-shell multi-TE data. SENSE (Pruessmann et al., 1999) served as the primary comparison with VC-MUSSELS (Bilgic et al., 2019) shown for appropriate scenarios (e.g., multi-TE data).

### 4.1. Experiment 1: HCP data

The retrospective acceleration experiment used publicly available multi-shell dMRI data from the HCP Lifespan project (Harms et al., 2018). The source data had 1.5 mm isotropic resolution, two diffusion shells at *b* = 1500 and 3000 s/mm^2^, 370 diffusion directions, 28 *b* = 0 volumes, nominal in-plane acceleration *R* = 1, and MB = 4. For each shell, 30 uniformly spread diffusion directions were selected by optimal one-to-one matching to a common 30-direction reference set, maximizing the total absolute cosine similarity between normalized b-vectors. Together with two *b* = 0 volumes, this yielded a 62-channel q-space-undersampled protocol. The k-space-fully-sampled reconstruction using the complete HCP q-space acquisition served as the reference. Complementary Cartesian masks at *R* = 3, 4, 5, 6, varying across neighboring diffusion directions, were applied to the synthetic multi-coil k-space. SENSE reconstructions provided the INR warm-up.

SENSE and DIFFUSE were evaluated against the k-space-fully-sampled reference reconstructed from the complete HCP q-space acquisition. Normalized mean squared error (NMSE) was computed for each axial slice *z*, diffusion direction *d*, and shell b-value *b* within a fixed, method-independent reference brain mask Ω:

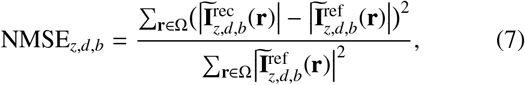

where 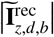 and 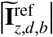 denote the reconstructed and reference motion-free magnitude DWIs, respectively.

Table 1 reports the reference-based NMSE at each acceleration factor. SENSE yielded lower mean NMSE at *R* = 3 and *R* = 4, while DIFFUSE yielded lower values at *R* = 5 and *R* = 6. DIFFUSE NMSE varied little across *R*, while SENSE NMSE increased sharply with acceleration. The q-space subset remained fixed at 30 directions per shell across acceleration factors, while the Fourier sampling density of each DWI varied with *R*. Figure 2 presents representative DWIs and residual maps, with the yellow numbers reporting mean NMSE across all slices. The fully sampled HCP reconstruction contains measurement noise, and closer reproduction of this noise contributes to the lower SENSE NMSE at *R* = 3 and *R* = 4. Across all acceleration factors, DIFFUSE produced less noisy images while preserving anatomical structure and yielded predominantly noise-like residuals. Coherent anatomical structure was visible in the SENSE residuals and became more pronounced at *R* = 5 and *R* = 6, particularly at *b* = 3000 s/mm^2^. These residual patterns highlight the advantage of the continuous shared signal field in DIFFUSE over slice-by-slice SENSE reconstruction. Conditioned on diffusion direction, b-value, and TE, the field couples available measurements so that common anatomical structure constrains each DWI. Shared parameterization reinforces signal components that remain consistent across spatial and acquisition coordinates and provides implicit regularization against incoherent noise.

**Table 1.**
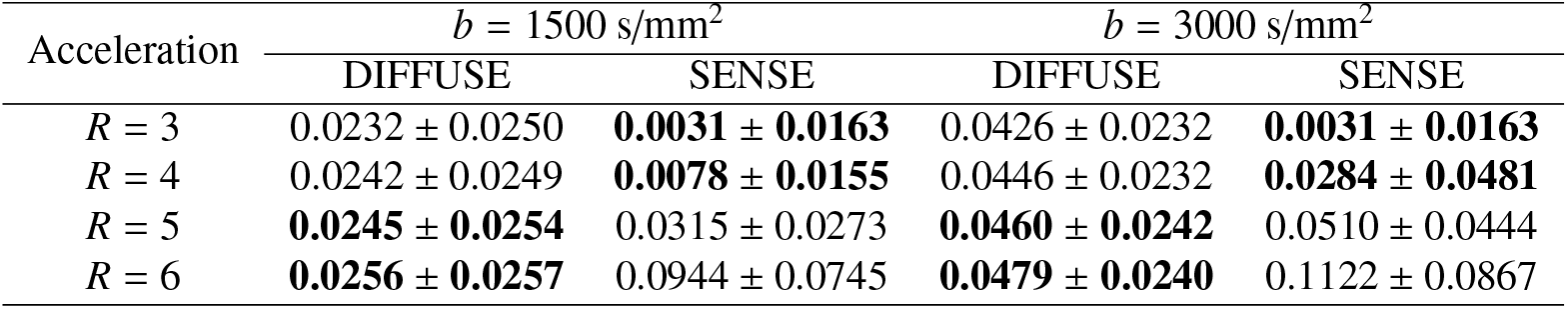
Reference-based NMSE for retrospective HCP multi-shell acceleration. Values are mean ± standard deviation (dimensionless) across valid axial slice–direction pairs, computed against the k-space-fully-sampled reference reconstructed from the complete HCP q-space acquisition.

| Acceleration | $b = 1500 \text{ s/mm}^2$ | | $b = 3000 \text{ s/mm}^2$ | |
| --- | --- | --- | --- | --- |
|  | DIFFUSE | SENSE | DIFFUSE | SENSE |
| $R = 3$ | $0.0232 \pm 0.0250$ | **$0.0031 \pm 0.0163$** | $0.0426 \pm 0.0232$ | **$0.0031 \pm 0.0163$** |
| $R = 4$ | $0.0242 \pm 0.0249$ | **$0.0078 \pm 0.0155$** | $0.0446 \pm 0.0232$ | **$0.0284 \pm 0.0481$** |
| $R = 5$ | **$0.0245 \pm 0.0254$** | $0.0315 \pm 0.0273$ | **$0.0460 \pm 0.0242$** | $0.0510 \pm 0.0444$ |
| $R = 6$ | **$0.0256 \pm 0.0257$** | $0.0944 \pm 0.0745$ | **$0.0479 \pm 0.0240$** | $0.1122 \pm 0.0867$ |

**Fig 2.**
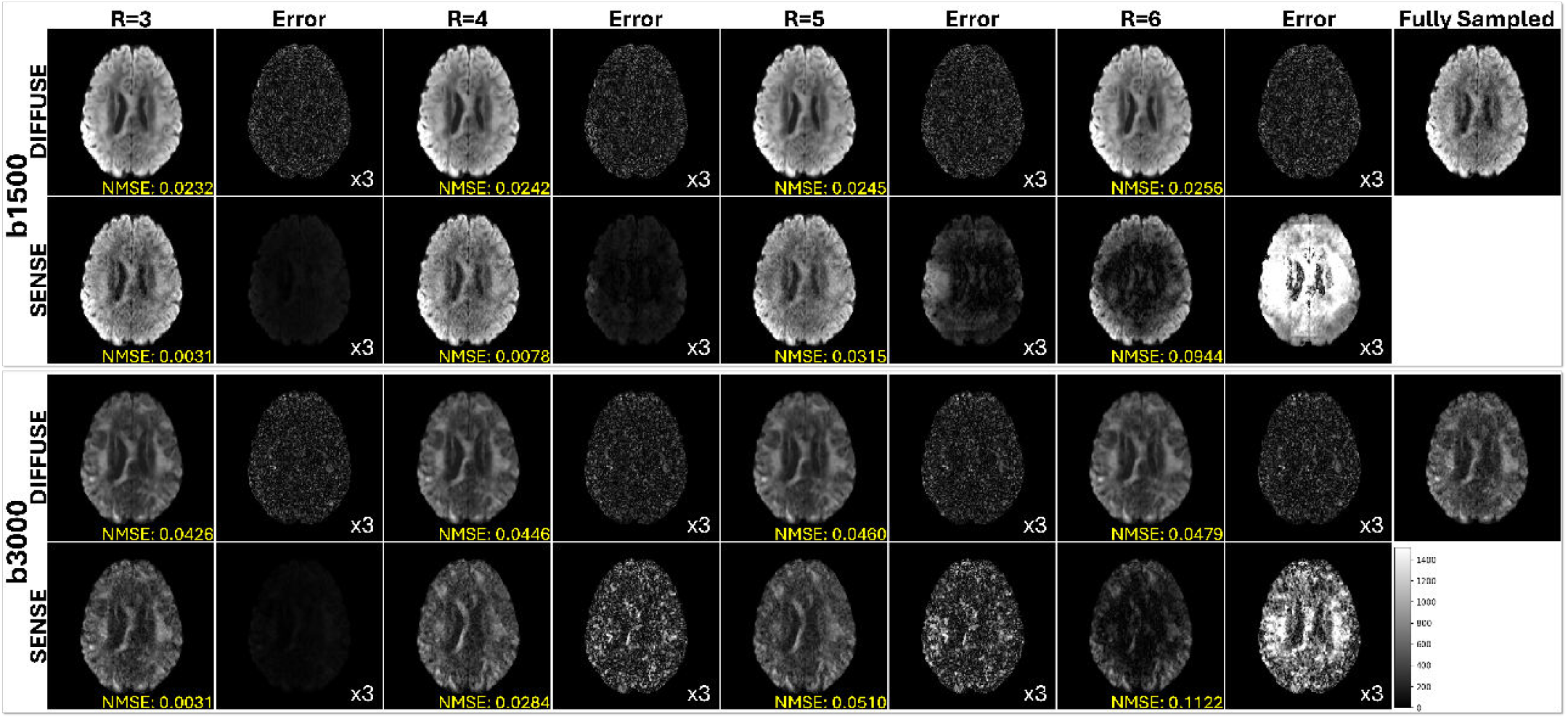
Retrospective HCP multi-shell acceleration. Diffusion-weighted images are shown for two shells (*b* = 1500 and 3000 s/mm^2^) under in-plane acceleration *R* = 3, 4, 5, and 6. For each acceleration factor, the SENSE and DIFFUSE outputs are shown alongside residual error maps computed against the k-space-fully-sampled reference reconstructed from the complete HCP q-space acquisition. Residual maps are displayed with three-fold intensity scaling. Yellow labels report the aggregate reference-based NMSE means from Table 1. The representative *R* = 6 panels show the largest visual separation, with stronger structured aliasing and residual signal for SENSE at *b* = 3000 s/mm^2^.

The DTI orientation maps in Figure 3 remained coherent for DIFFUSE from *R* = 3 to *R* = 6 in both shells. SENSE developed marked directional disruption at high acceleration, most clearly at *R* = 6. DIFFUSE also retained smooth, anatomically organized contrast in the corresponding FA and MD maps (Supplementary Figures S1 and S2), while SENSE exhibited spatially structured intensity variation at high *R*. The DIFFUSE ODF fields in Figure 4 retained consistent large-scale organization and local glyph alignment across all acceleration factors and shells, thereby indicating that despite the larger NMSE at lower acceleration factors, no information is lost (and only noise was removed).

**Fig 3.**
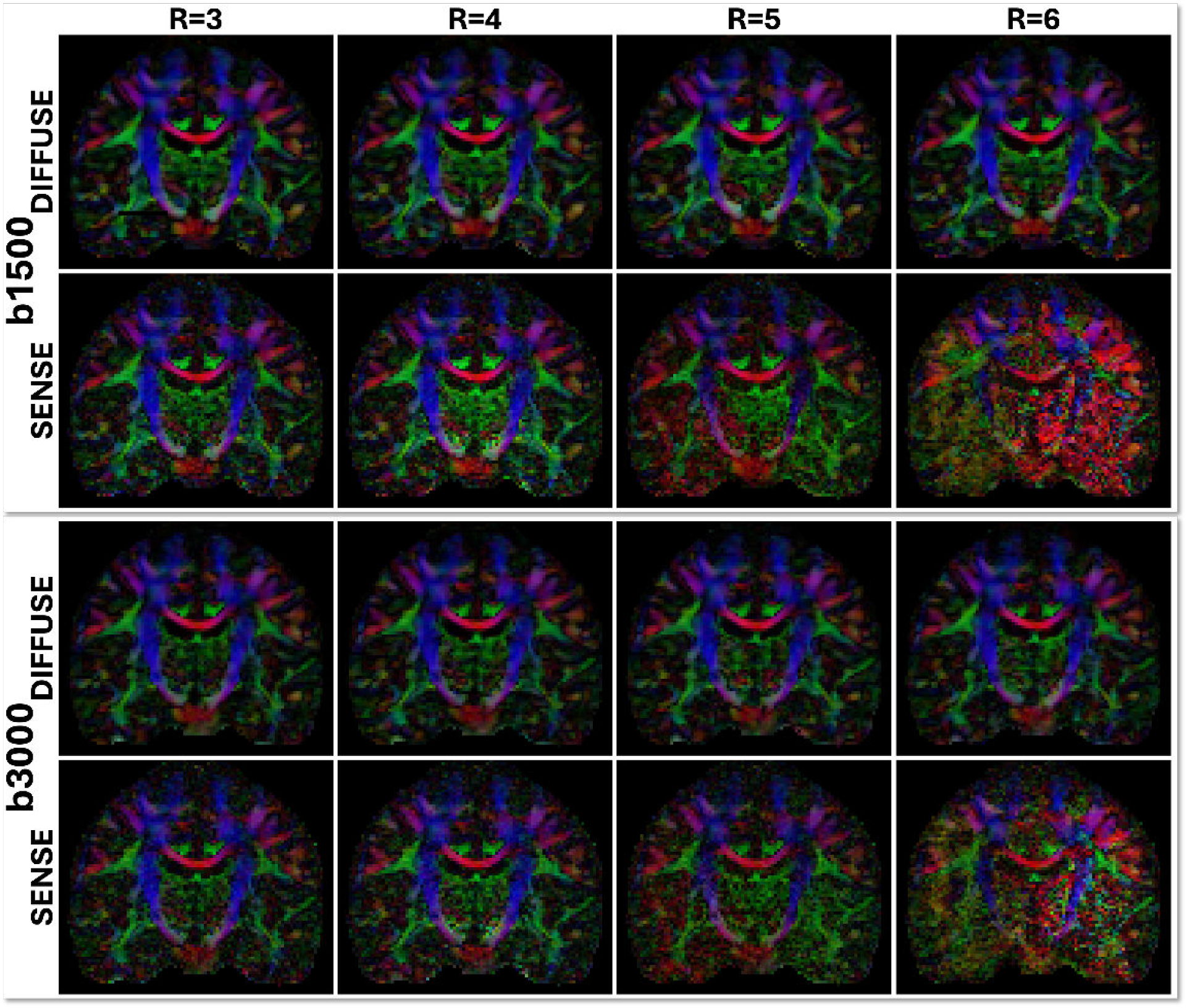
DTI orientation maps for retrospective HCP multi-shell acceleration. Color-encoded DTI orientation maps from SENSE and DIFFUSE are shown for the same shells and acceleration factors as in Figure 2. These panels provide a qualitative comparison of DTI orientation maps across the displayed acceleration settings.

**Fig 4.**
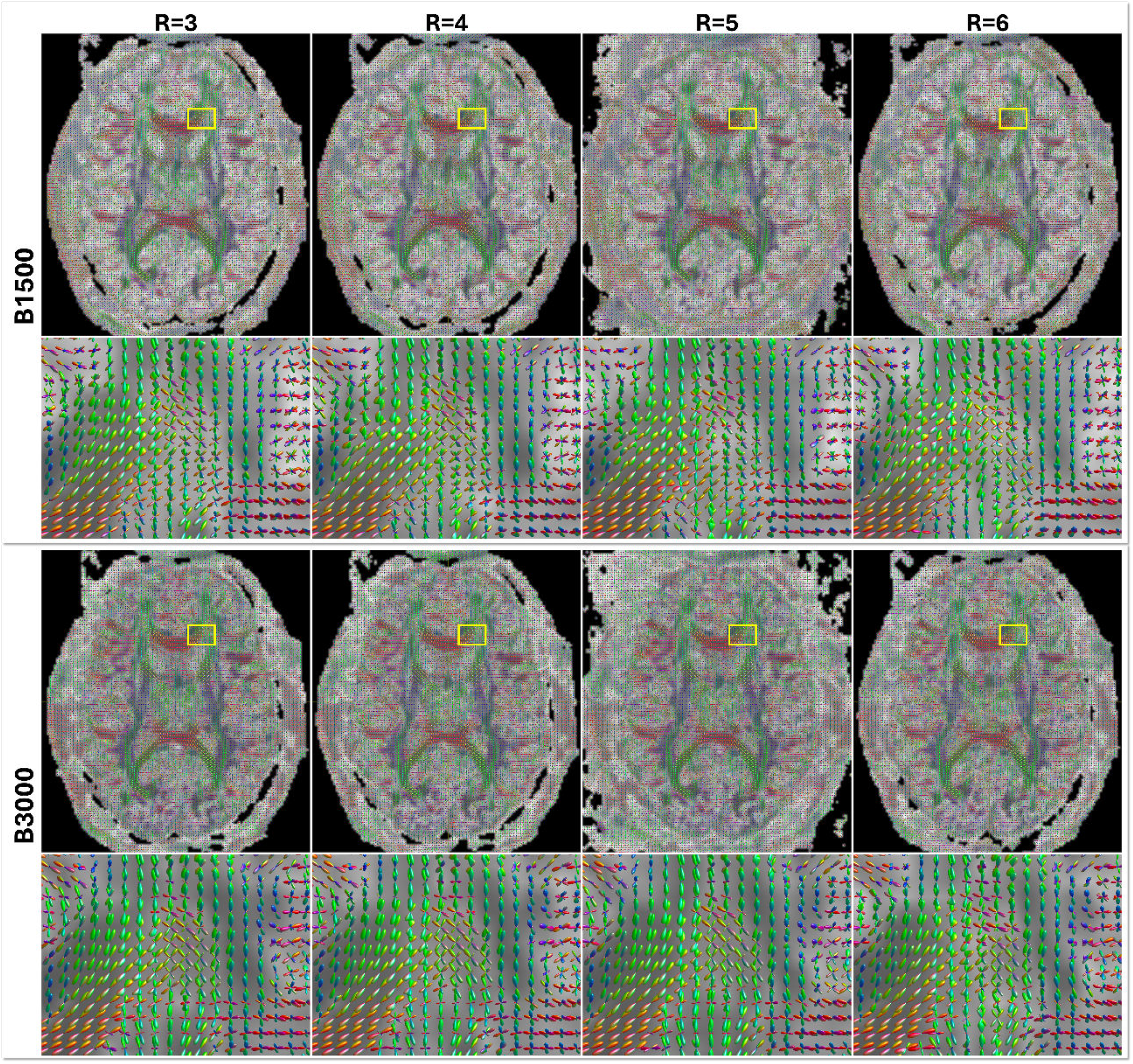
ODF fields derived from DIFFUSE for retrospective HCP multi-shell acceleration without motion. Columns show *R* = 3, 4, 5, and 6, while the upper and lower panel groups show *b* = 1500 and 3000 s/mm^2^, respectively. Each shell includes a whole-slice ODF field and a magnified view of the yellow region of interest. The panels provide a within-method qualitative view across acceleration factors using a fixed orientation and color convention.

### 4.2. Experiment 2: Motion-aware spatial mapping with multiple shots

In this experiment, rigid head motion was simulated with different motion applied to each diffusion direction. Additionally, assuming multi-shot acquisition, a typical scenario where head motion correction is imperative during reconstruction, different motion parameters were used to simulate shot-to-shot head motion. Each of the two shots was undersampled at *R* = 6. For each diffusion-weighted volume, a prescribed rigid motion was applied to the 3D volume with rotation of up to 3 degrees (keeping center of the image as the origin) and translation of 2 to 5 pixels. The two shot motion trajectories are shown in Figure 5. ANTsPy rigid registration estimated the transform from each SENSE estimated volume with the corresponding shot mean-*b*0 image as the reference. The estimated transforms were used during the INR fitting. These transforms were found to be adequate for removing the motion artifacts. However, the framework is general enough to allow iterative updates to the motion parameters during fitting to remove any additional motion related blurring.

**Fig 5.**
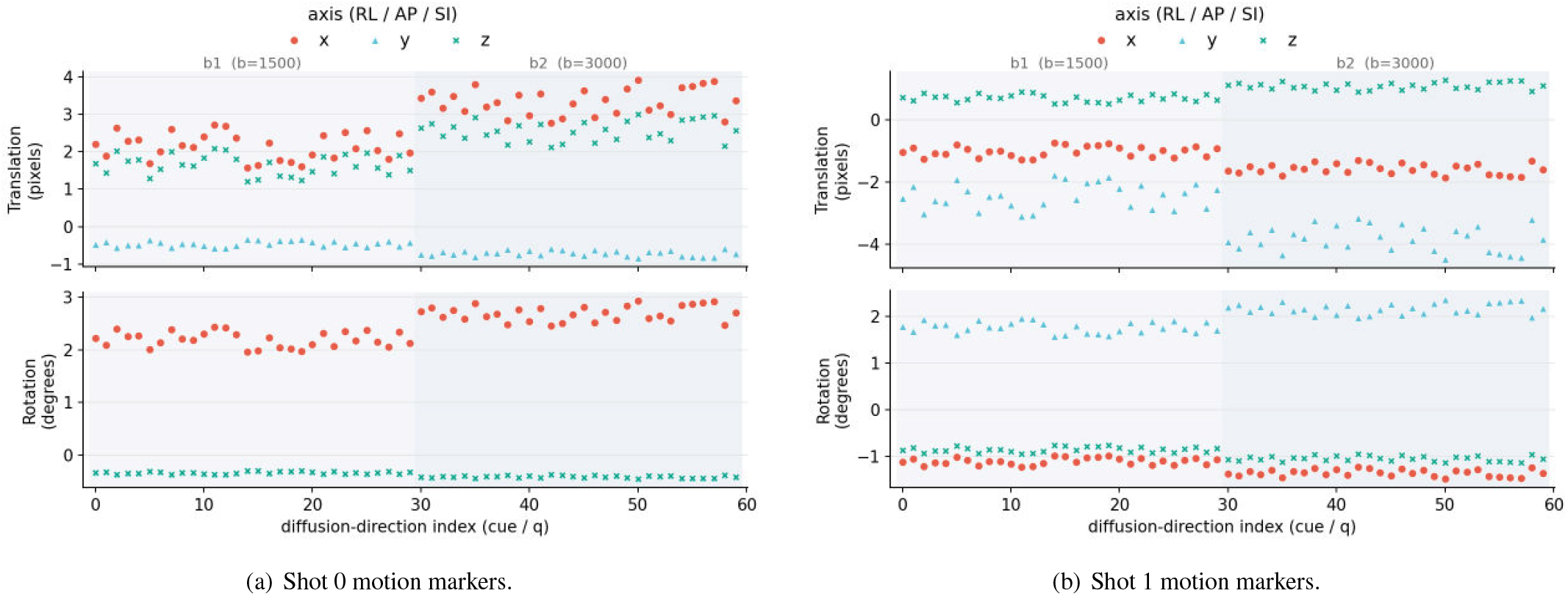
Controlled HCP inter-shot motion simulation. Translation and rotation markers are shown for the two simulated shots used in the HCP motion stress test, with each shot undersampled at *R* = 6. Markers are indexed by diffusion direction, with shaded regions corresponding to the *b* = 1500 and 3000 s/mm^2^ shells. The *b* = 0 measurements remain stationary. Across the 60 diffusion-weighted directions, translation-vector norms increase from 2 to 5 image-grid pixels (voxel-index units) and rotation-vector norms from 2^°^ to 3^°^. The two shot trajectories were generated with seeds 42 and 43 and define the rigid poses used to synthesize motion-corrupted k-space.

A consistent ranking was observed across both shells in Table 2. Motion-aware DIFFUSE achieved the lowest mean NMSE, followed by static DIFFUSE, SENSE, and VC-MUSSELS. Relative to static DIFFUSE, motion-aware spatial mapping and b-vector reorientation reduced mean NMSE by 72% at *b* = 1500 s/mm^2^ and 66% at *b* = 3000 s/mm^2^. The reconstructed slices and residual maps in Figure 6 reflect this ranking. Motion-aware DIFFUSE preserved the fully sampled anatomy most closely and produced the weakest structured residuals in both shells. Static DIFFUSE retained coherent residual anatomy, whereas SENSE and VC-MUSSELS displayed more extensive residual signal and image distortion. The motion-aware DIFFUSE ODF fields in Figure 7 retained coherent whole-slice organization in both shells and locally ordered orientations in the magnified regions.

**Table 2.** Reference-based NMSE for controlled HCP inter-shot motion reconstruction. Values are mean ± standard deviation (dimensionless) across valid axial slice–direction pairs, computed against the motion-free HCP reference. The VC-MUSSELS results are also shown for comparison.

| Method | $b = 1500$ s/mm<sup>2</sup> | $b = 3000$ s/mm<sup>2</sup> |
| --- | --- | --- |
| Motion-aware |  |  |
| DIFFUSE | **0.050 ± 0.028** | **0.077 ± 0.032** |
| Static |  |  |
| DIFFUSE | 0.181 ± 0.123 | 0.227 ± 0.108 |
| SENSE | 0.240 ± 0.120 | 0.297 ± 0.125 |
| VC-MUSSELS | 0.719 ± 0.185 | 0.741 ± 0.228 |

**Fig 6.**
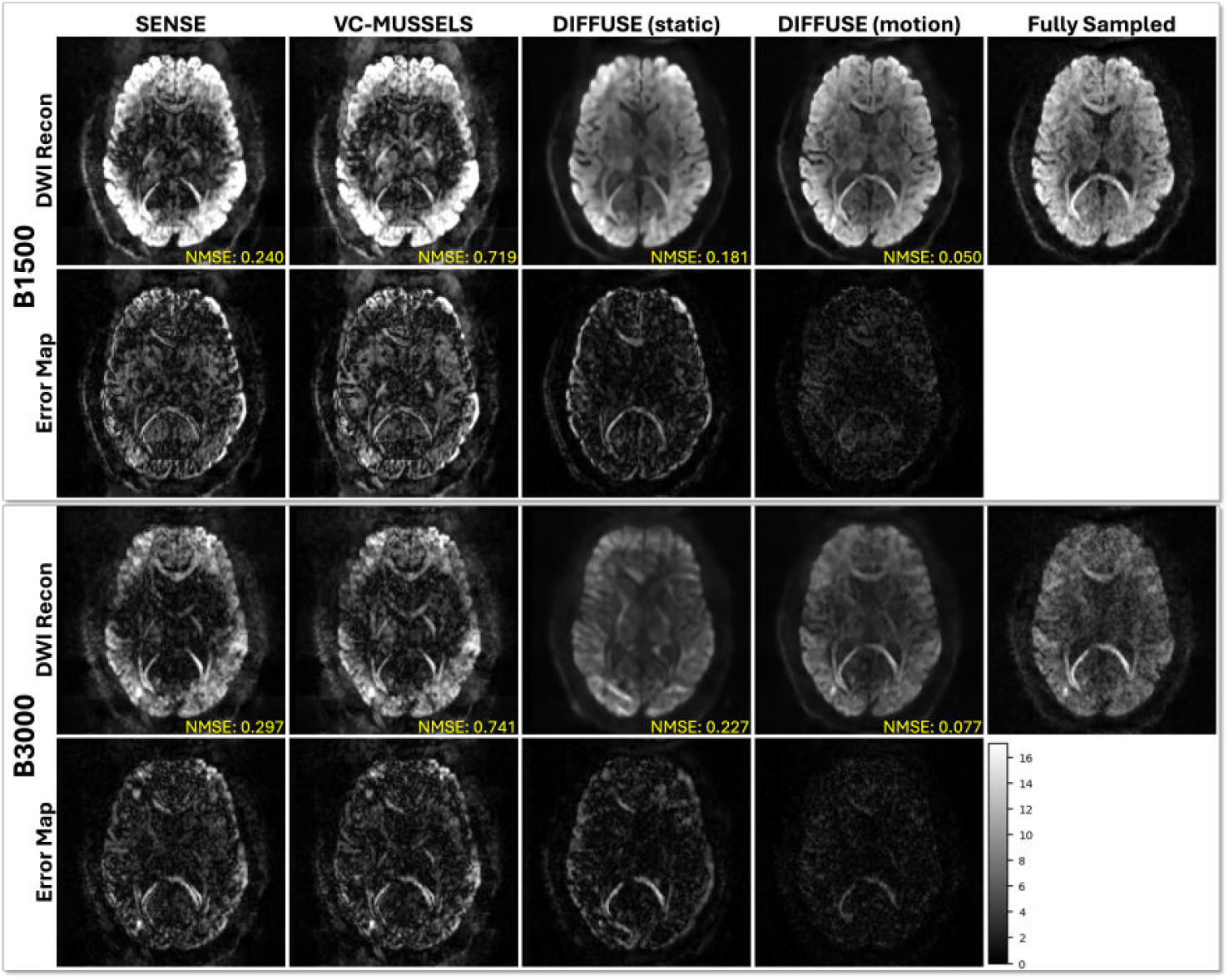
Controlled HCP inter-shot motion stress test at *R* = 6. Each shot is undersampled at *R* = 6. Diffusion-weighted images and residual maps are shown for two shells (*b* = 1500 and 3000 s/mm^2^). Columns compare SENSE, VC-MUSSELS, static DIFFUSE, motion-aware DIFFUSE, and the k-space-fully-sampled motion-free reference reconstructed from the complete HCP q-space acquisition. Yellow labels report the aggregate reference-based NMSE means from Table 2.

**Fig 7.**
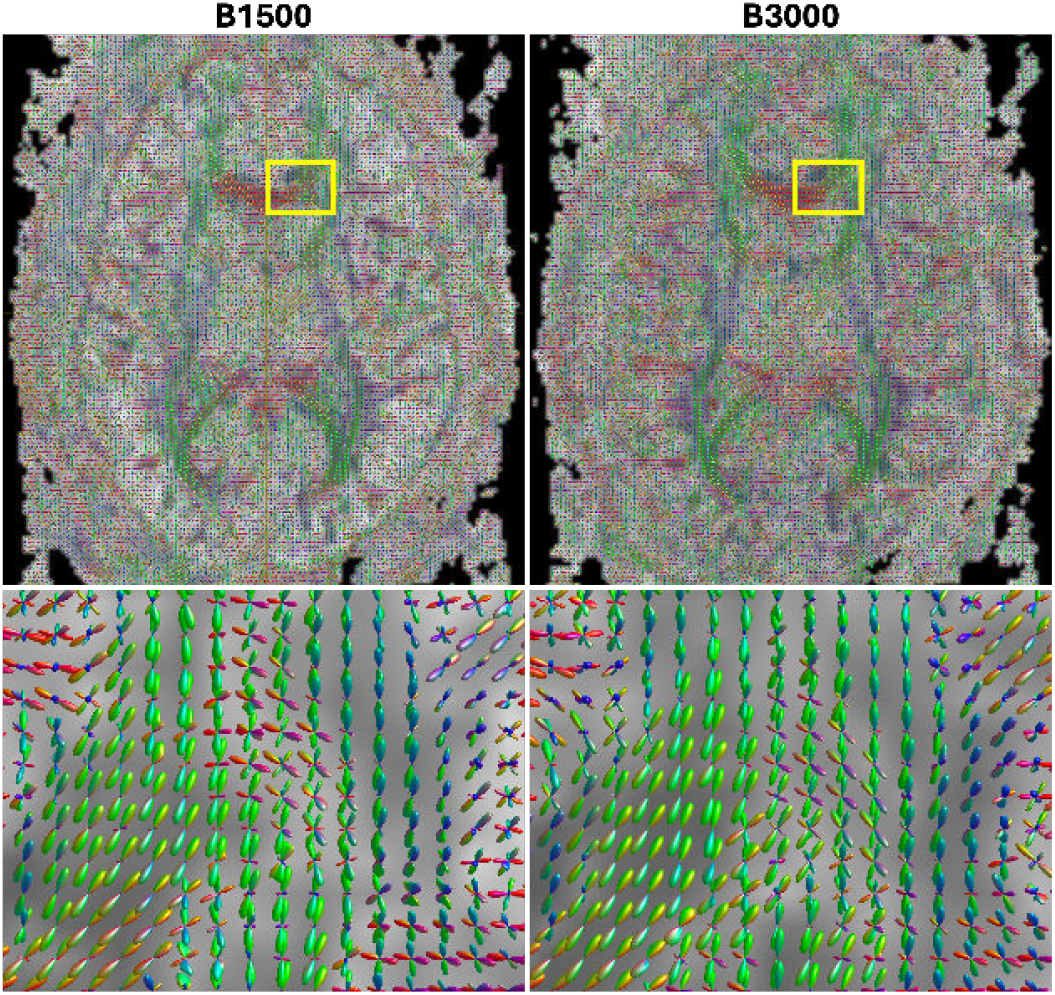
ODF visualizations after motion-aware reconstruction in the controlled HCP inter-shot motion experiment at *R* = 6. The left and right columns show *b* = 1500 and 3000 s/mm^2^, respectively. The upper row shows whole-slice ODF fields, and the lower row magnifies the yellow regions of interest. The fields provide a qualitative within-method visualization under a fixed orientation and color convention.

### 4.3. Experiment 3: Prospective 1.3 mm at undersampling of R = 2, 4, 5

The prospective experiment comprised three separately acquired scans at acceleration factors of *R* = 2, 4, 5 on a Siemens Prisma scanner. The common nominal protocol used 1.3 mm isotropic resolution, 6*/*8^*th*^ partial Fourier, and MB = 2. The shells at *b* = 1500 and 3000 s/mm^2^ each used an undersampled q-space protocol containing 30 diffusion directions. Each forward-encoded scan was paired with a PA acquisition containing six reverse phase-encoded volumes for topup-based distortion correction. Registration of SENSE or initial INR images supplied the fixed transforms used for motion-aware reconstruction. Each acceleration setting was fitted separately. The scans were separate acquisitions without a common fully sampled reference. Accordingly, the evaluation was limited to qualitative comparisons within each acquisition and did not test the reference-based NMSE crossover observed in the retrospective HCP study.

Within each prospective acquisition, the DWIs in Figure 8 exhibited robust reconstruction with DIFFUSE than with SENSE or VC-MUSSELS. The visual separation is most pronounced at *b* = 3000 s/mm^2^. In Figure 9, the DTI orientation maps and FA maps from DIFFUSE contained less speckled noise than the SENSE maps, while the DIFFUSE MD maps showed more continuous tissue contrast. The ODF fields in Figure 10 are more spatially ordered for DIFFUSE than for SENSE in both the whole-slice and magnified views. These within-acquisition observations support the feasibility of prospective 1.3 mm multi-shell reconstruction at the three acquired acceleration settings.

**Fig 8.**
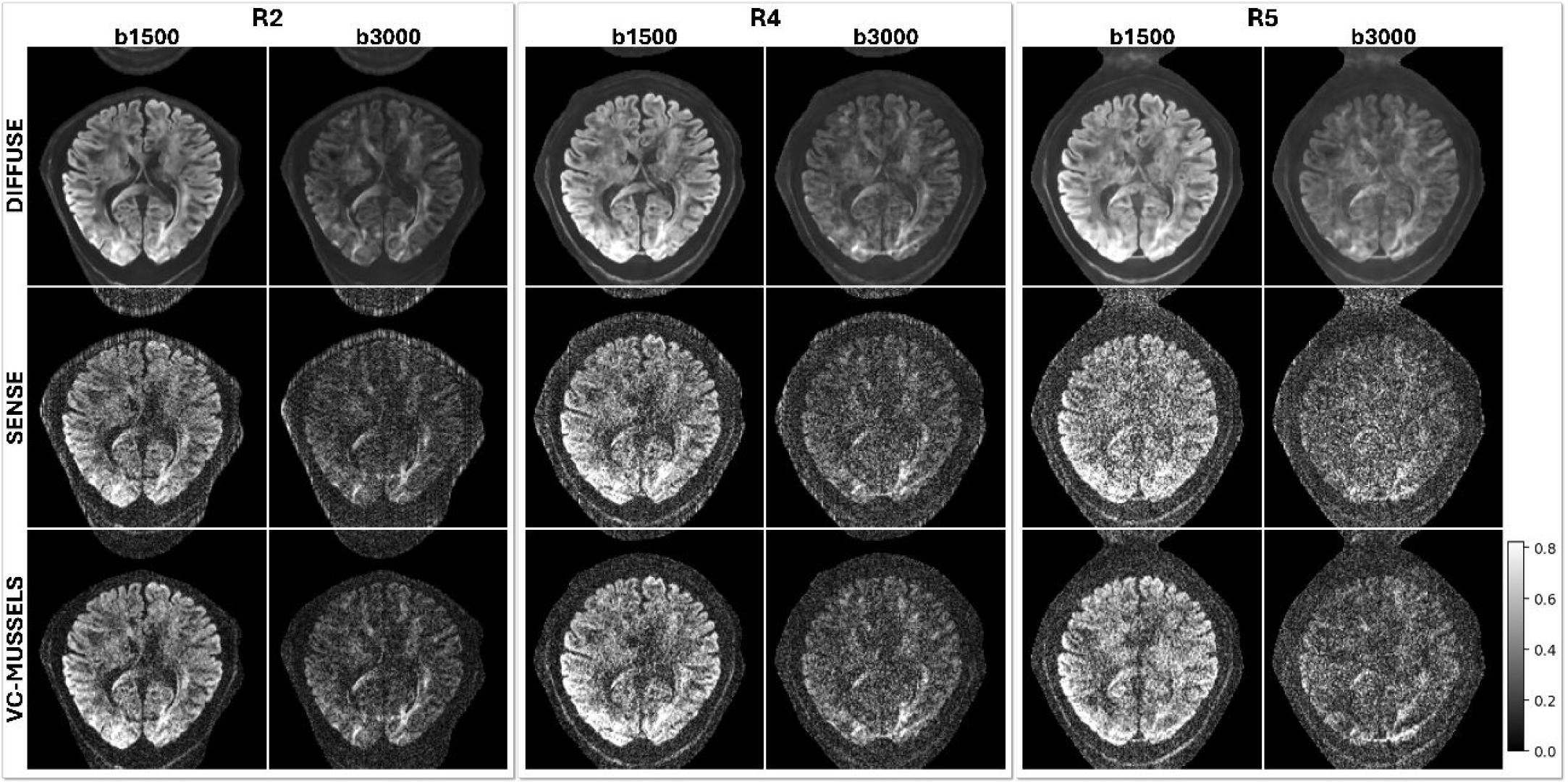
Prospective high-resolution in vivo multi-shell reconstruction. Diffusion-weighted images are shown for *R* = 2, 4, 5 with MB = 2 at *b* = 1500 and 3000 s/mm^2^. For each acceleration factor and shell, DIFFUSE, SENSE, and VC-MUSSELS are compared on the prospectively acquired high-resolution data. The three settings are separate acquisitions, with *R* = 2 serving as a lower-acceleration comparator.

**Fig 9.**
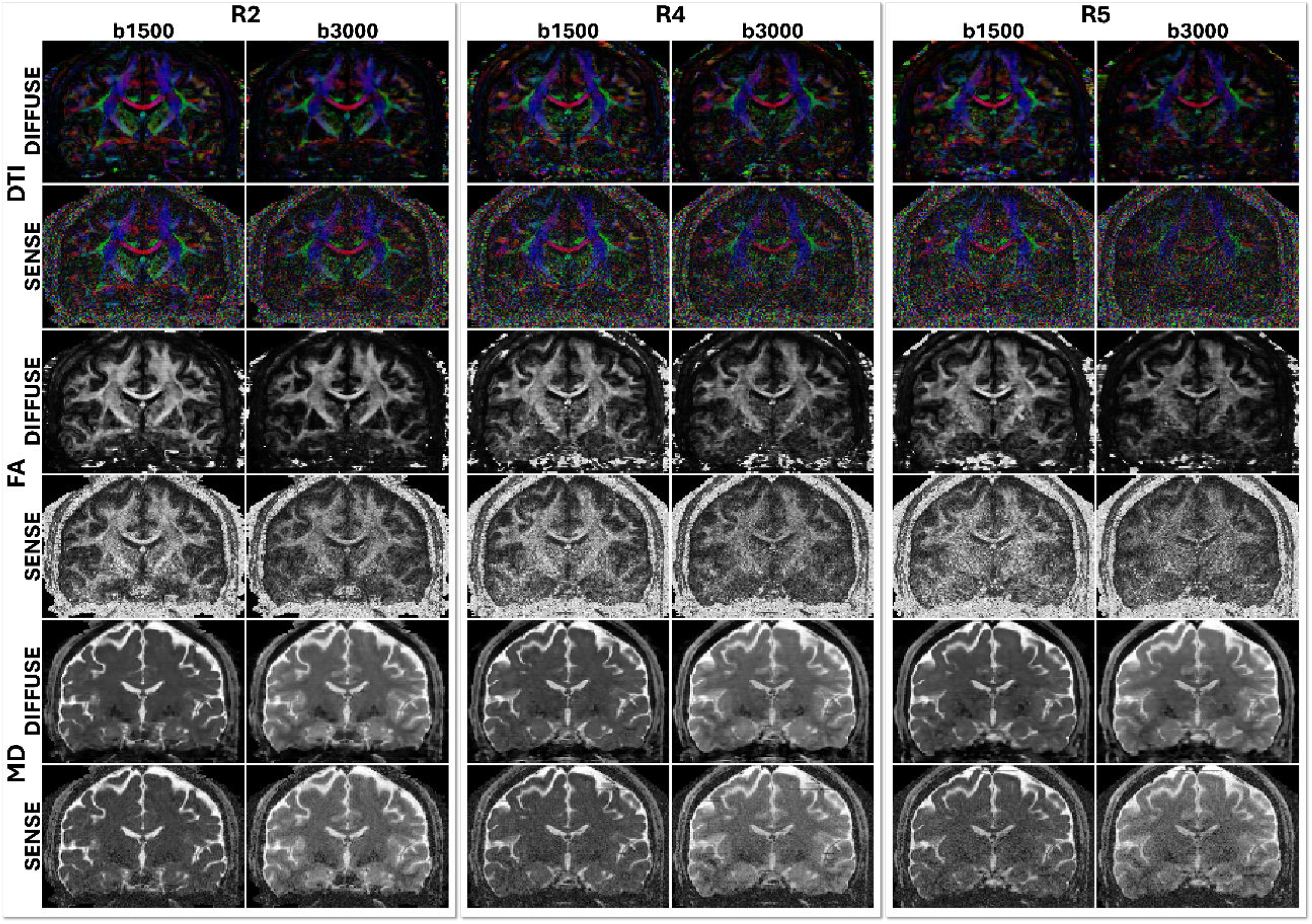
Tensor-derived maps for the prospective 1.3 mm isotropic acquisition. Columns follow *R* = 2, 4, 5 and the two shells (*b* = 1500 and 3000 s/mm^2^). Rows show color-encoded DTI orientation maps, FA maps, and MD maps for DIFFUSE and SENSE. The panels provide reference-free qualitative comparisons of the DTI orientation maps, FA maps, and MD maps.

**Fig 10.**
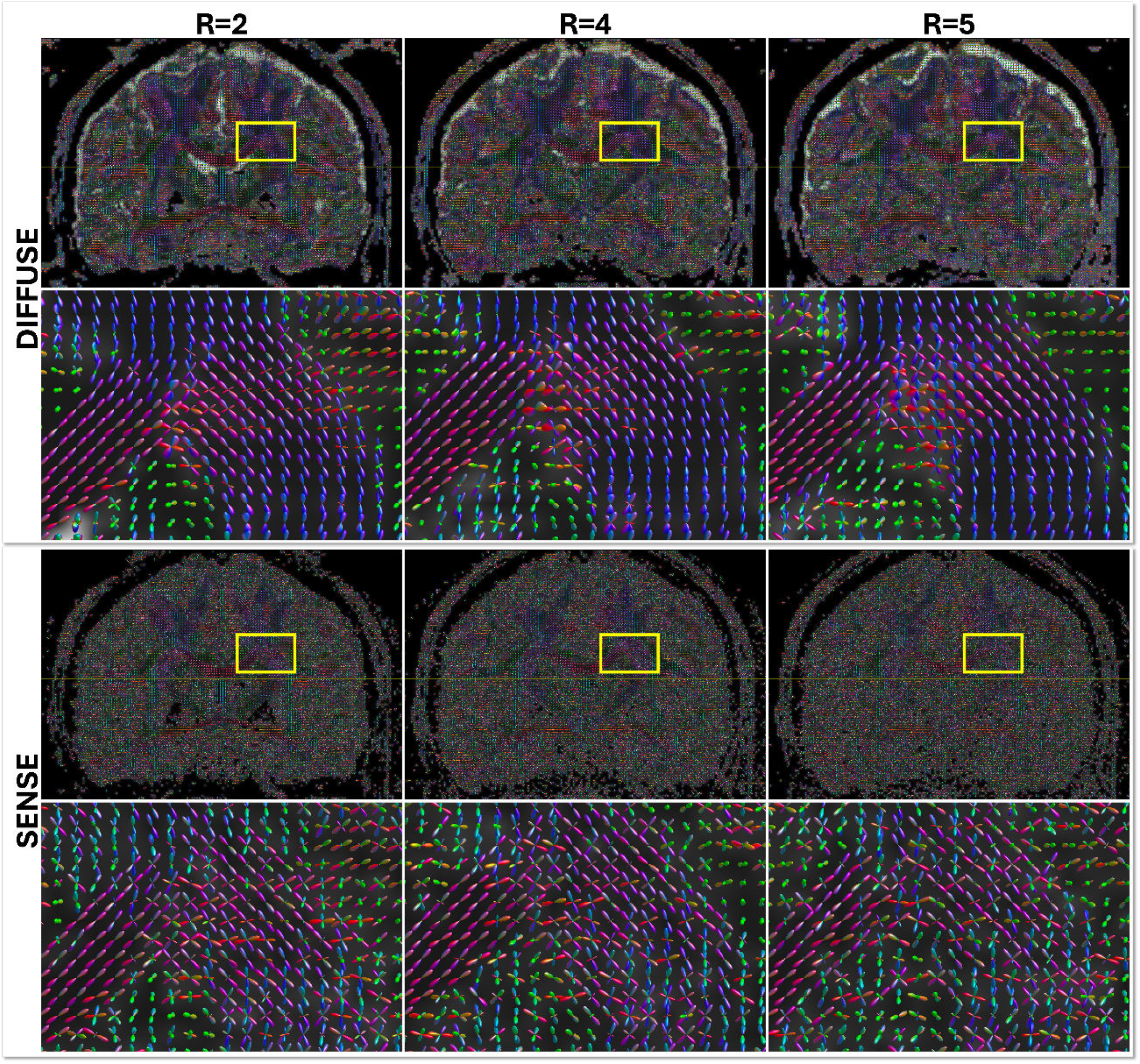
ODF fields from DIFFUSE and SENSE for the prospective 1.3 mm isotropic multi-shell acquisition. Columns show reconstructions at *R* = 2, 4, and 5 with MB = 2. For each method, the upper panels present whole-slice ODF fields and the lower panels magnify the yellow regions of interest. The *R* = 2 acquisition provides a lower-acceleration comparator, and all panels support qualitative reference-free comparison within each acquisition.

### 4.4. Experiment 4: One shared field represents multi-b-value multi-TE data

The multi-shell multi-TE experiment used a time-division multiplexed EPI acquisition (Ji et al., 2022a; Liu et al., 2024). The protocol combined *b* = 1000, 2000, and 3000 s/mm^2^ with TE values of 79, 109, and 139 ms. Each of the nine b-value/TE combinations used an undersampled q-space protocol containing 32 diffusion directions and one *b* = 0 volume at 2.0 mm isotropic resolution. In-plane acceleration was *R* = 3, with complementary *k*_*y*_ shifts across neighboring diffusion directions and echoes. DIFFUSE used the same five-dimensional INR formulation as in the preceding experiments to jointly reconstruct the complete acquired grid of three b-values and three TEs.

As shown in Figure 11, a single fitted field generated outputs for all nine acquired conditions, establishing joint-representation capability. Across the complete 3 × 3 grid, DIFFUSE maintained spatially continuous anatomy, whereas SENSE and VC-MUSSELS displayed more granular within-brain signal. The common intensity scale retained condition-dependent contrast across b-values and TEs, including lower signal in the *b* = 2000 and *b* = 3000 s/mm^2^ rows. The ODF fields in Figure 12 retained spatial organization across all nine conditions, and each magnified region exhibited locally ordered orientations. Corresponding coronal FA, MD, and DTI orientation maps across the complete 3 × 3 b-value/TE grid are provided in Supplementary Figures S3–S5. Together, the DWI and ODF observations document condition-dependent signal contrast and directional structure throughout the joint reconstruction.

**Fig 11.**
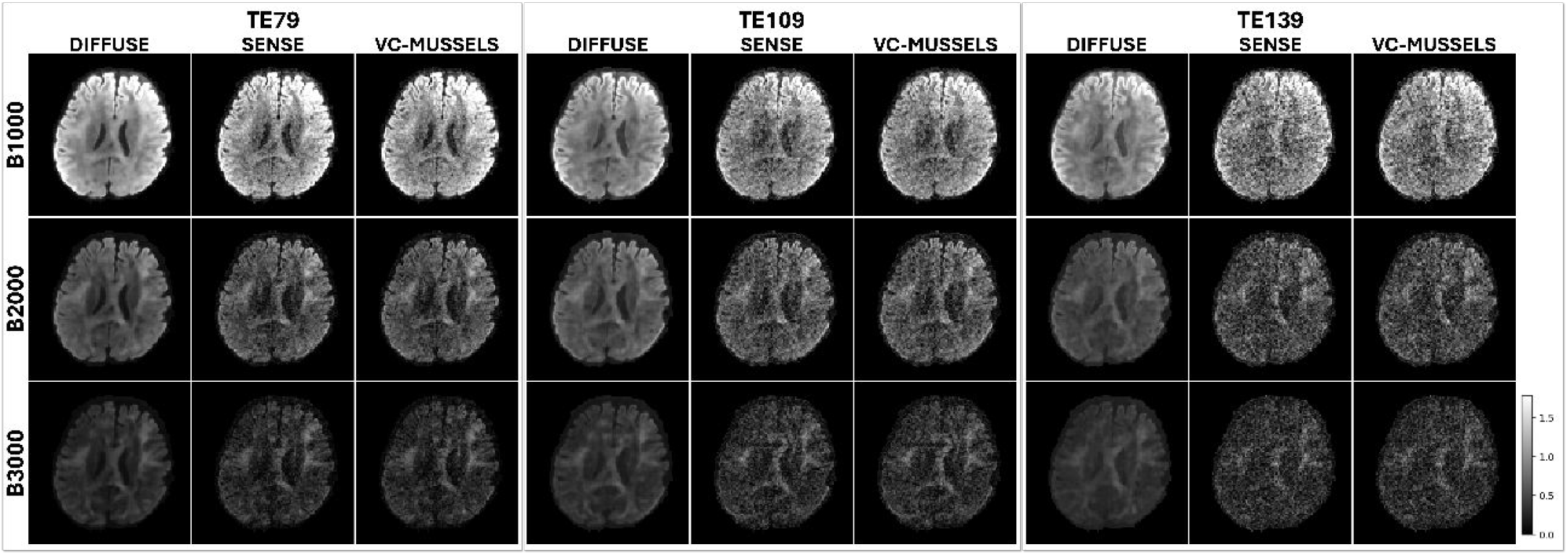
In vivo multi-shell multi-TE reconstruction. Diffusion-weighted images at *b* = 1000, 2000, and 3000 s/mm^2^ are shown for TE = 79, 109, and 139 ms. For each b-value/TE combination, DIFFUSE, SENSE, and VC-MUSSELS are compared under complementary *k*_*y*_-shifted sampling across neighboring directions and echoes (Ji et al., 2022a; Liu et al., 2024). SENSE was matched to the DIFFUSE brain mean within each b-value/TE condition, and all panels were displayed with one global intensity scale. Corresponding FA maps are provided in Supplementary Figure S3.

**Fig 12.**
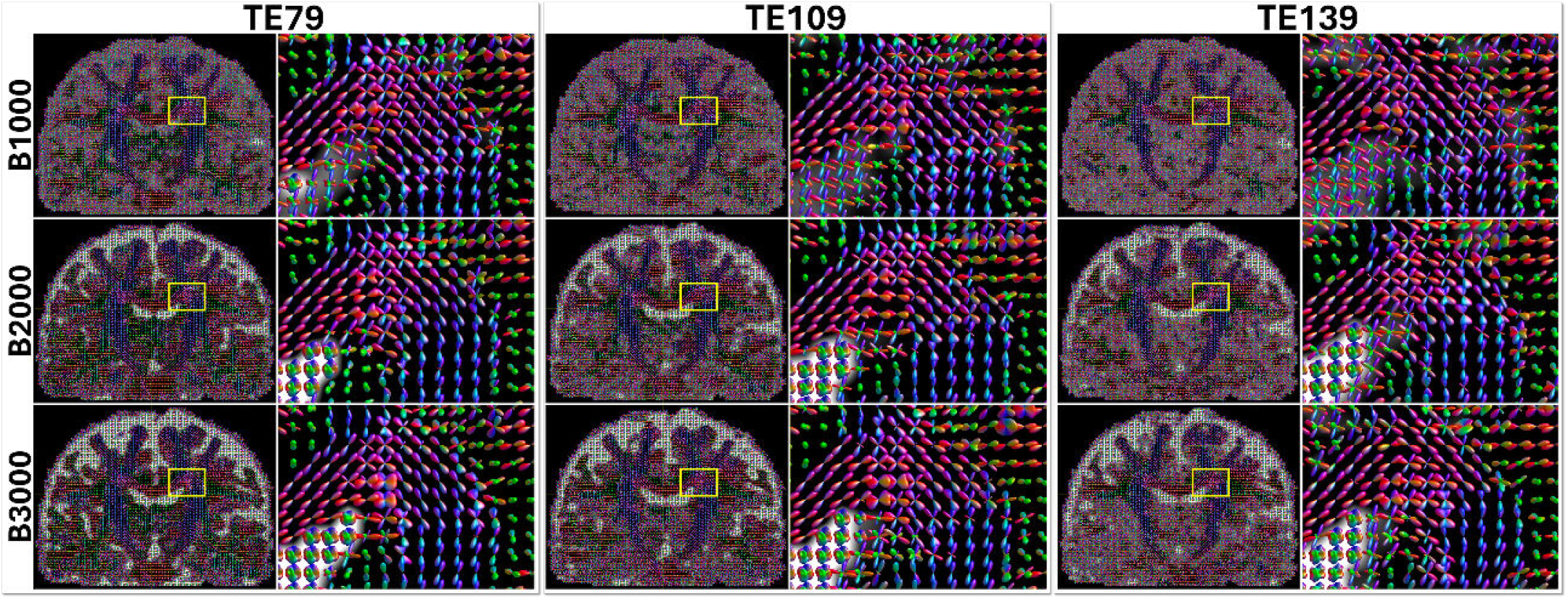
ODF fields derived from DIFFUSE across the nine TDM-EPI b-value/TE combinations. Rows show *b* = 1000, 2000, and 3000 s/mm^2^, while columns show TE = 79, 109, and 139 ms. Each combination pairs a whole-slice ODF field with a magnified view of the yellow region of interest. The grid provides a qualitative display of representation capacity across the nine acquired conditions.

## 5. Discussion

Across the evaluated settings, the image comparisons and available residual maps consistently favored DIFFUSE in recovering the underlying diffusion-weighted image structure. As acceleration increased, fewer k-space samples remained available for each DWI, making independent reconstruction increasingly ill-conditioned. DIFFUSE uses complementary samples from related diffusion directions and shells to constrain missing k-space locations. Its advantage therefore became most apparent at high acceleration. At lower acceleration, the lower reference NMSE of SENSE may partly reflect closer reproduction of noise in the HCP reference. DIFFUSE instead empha-sizes anatomical structure supported across related measurements. The shared signal field reinforces anatomical structure that recurs across measurements, while measurement-specific forward operators preserve each acquisition’s contrast and encoding. Each DWI therefore retains its genuine contrast, while cross-measurement information contributes more as independent reconstruction becomes less constrained.

The motion experiment showed why spatial and angular correspondence matters for cross-measurement sharing. Reference-space mapping and b-vector reorientation place measurements acquired at different poses in a common spatial and angular frame. This combined alignment explains the improvement of motion-aware DIFFUSE over static DIFFUSE. The prospective 1.3 mm and TDM-EPI experiments further showed that DIFFUSE can operate directly on acquired high-resolution multi-shell and multi-b/multi-TE data. Because the signal field is fitted anew for each examination, it adapts to the measured anatomy and acquisition while using complementary information across directions, shells, and TEs. Measurement-specific encoding remains in the forward model, so genuine signal differences associated with b-value and TE are preserved. The resulting DWIs, tensor-derived maps, and ODFs remain compatible with standard downstream diffusion analyses. In practice, this approach makes fuller use of related measurements within a fixed acquisition budget and supports protocols spanning richer combinations of directions, shells, and TEs.

Subject-specific fitting avoids population-level training but remains computationally intensive for each examination. Table 3 reports INR fitting times and peak GPU memory for the prospective 1.3 mm and multi-*b*-value multi-TE experiments. The reported costs jointly reflect the volume size and fitted *b*-value/TE grid. During the prospective 1.3 mm reconstructions, the appropriate stopping point varied with acceleration. Prolonged optimization may increasingly fit measurement noise in more ill-conditioned high-acceleration settings, whereas premature stopping can leave the reconstruction overly smooth. Future work could develop acceleration-aware stopping criteria that balance detail recovery against noise fitting. Our current optimization strategy uses SENSE-based warm-up, which not only reduces computational burden but also provides a “ghost-free” or unaliased starting point, thereby improving optimization robustness. Another direction is to jointly or alternately refine the signal field, phase maps, rigid-pose parameters, and nonrigid deformation fields within the measurement-specific forward model. This extension could coordinate tissue representation and acquisition effects within the same data-consistency formulation.

**Table 3.** INR fitting time and peak GPU memory for the 32-coil prospective 1.3 mm and multi-*b*-value multi-TE experiments. Voxel sizes are isotropic, and fitting times were measured on NVIDIA RTX A6000 GPUs (48 GB).

| Experiment | Voxel size | Volume size (voxels) | Fitted $b$-value/TE grid | INR fitting time | Peak memory |
| --- | --- | --- | --- | --- | --- |
| Prospective 1.3 mm | 1.3 mm | $170 \times 170 \times 100$ | 2 shells × 1 TE | 46.6 hours | 12.2 GB |
| Multi-$b$-value multi-TE | 2.0 mm | $100 \times 100 \times 60$ | 3 shells × 3 TEs | 36.0 hours | 20.3 GB |

## 6. Conclusion

We presented DIFFUSE, a subject-specific joint k–q–TE reconstruction framework that fits a five-dimensional diffusion-signal field directly to complementary undersampled multi-coil k-space measurements across diffusion encodings and echo times. DIFFUSE maintained stable NMSE as the acceleration factor increased and achieved 57–73% lower NMSE than SENSE at *R* = 6. At the same acceleration, motion-aware DIFFUSE achieved 74–79% lower NMSE than SENSE. The prospective study yielded 1.3 mm isotropic multi-shell reconstructions, and a single joint fit reconstructed multi-shell, multi-TE data spanning three b-values and three echo times. Together, the experiments show that a single subject-specific model can jointly reconstruct multi-shell, multi-TE diffusion signals from complementary undersampled multi-coil k-space measurements.

## Acknowledgements

This work was supported by the National Institute of Mental Health of the National Institutes of Health under Award Number R01MH116173. The content is solely the responsibility of the authors and does not necessarily represent the official views of the National Institutes of Health.

## Declaration of generative AI and AI-assisted technologies in the manuscript preparation process

During the preparation of this work the authors used Claude (Anthropic) in order to improve the language and readability of the manuscript. After using this tool, the authors reviewed and edited the content as needed and take full responsibility for the content of the published article.

