## Supplementary Material for "Joint reconstruction of multi-contrast relaxation-diffusion imaging"

Supplementary Figures S1 and S2 provide the FA and MD maps for the retrospective HCP multi-shell acceleration experiment. Supplementary Figures S3–S5 provide FA, MD, and DTI orientation maps at coronal slice $y = 53$ across the complete $3 \times 3$ b-value/TE grid in the multi-shell multi-TE experiment.

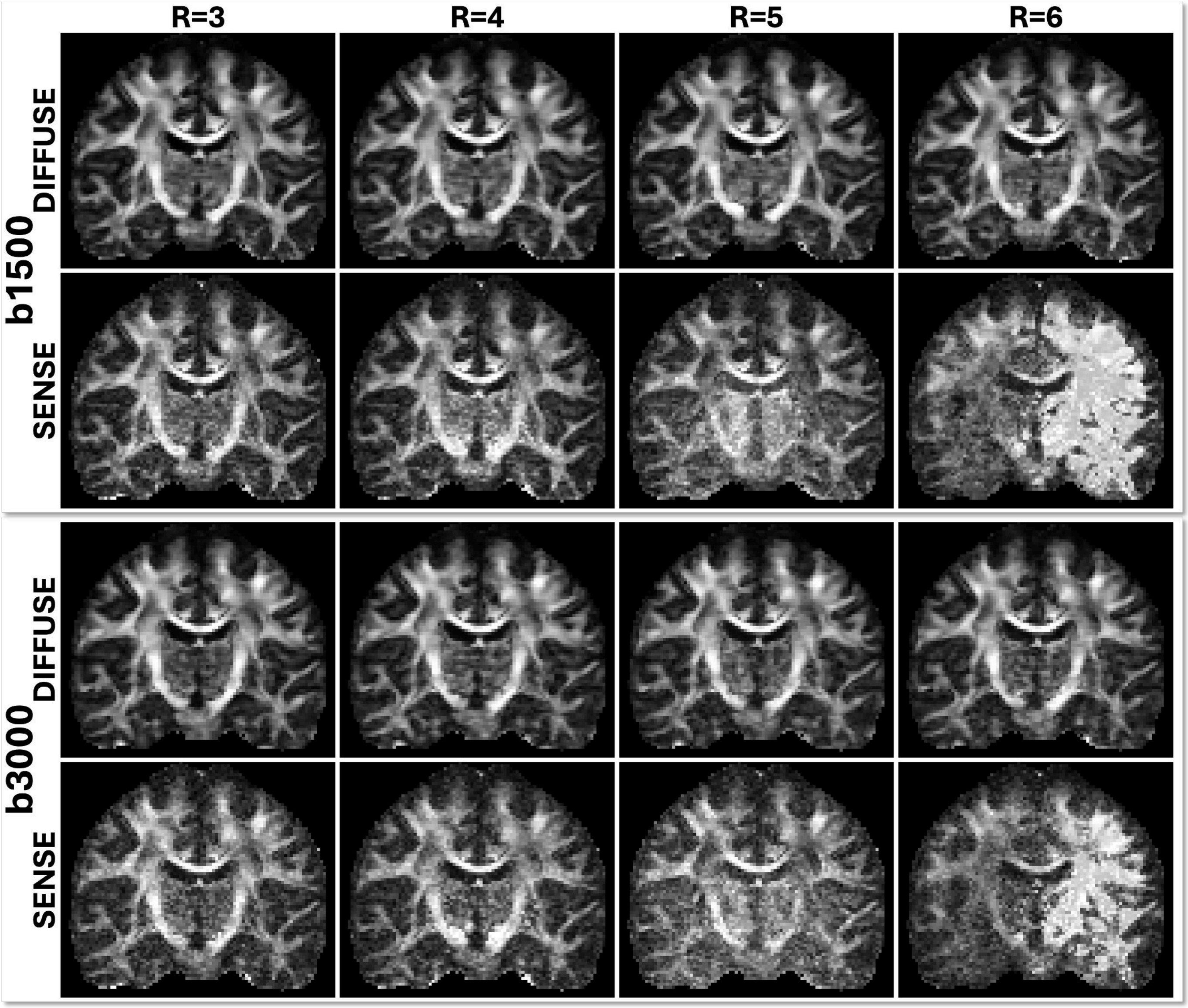


**Fig. S1. Fractional anisotropy (FA, scale 0–1) for retrospective HCP multi-shell acceleration. Maps are arranged by acceleration factor and shell, with SENSE and DIFFUSE compared under matched tensor-estimation settings. DIFFUSE reduces visible aliasing-driven contamination of FA at $R = 5$ and $R = 6$.**

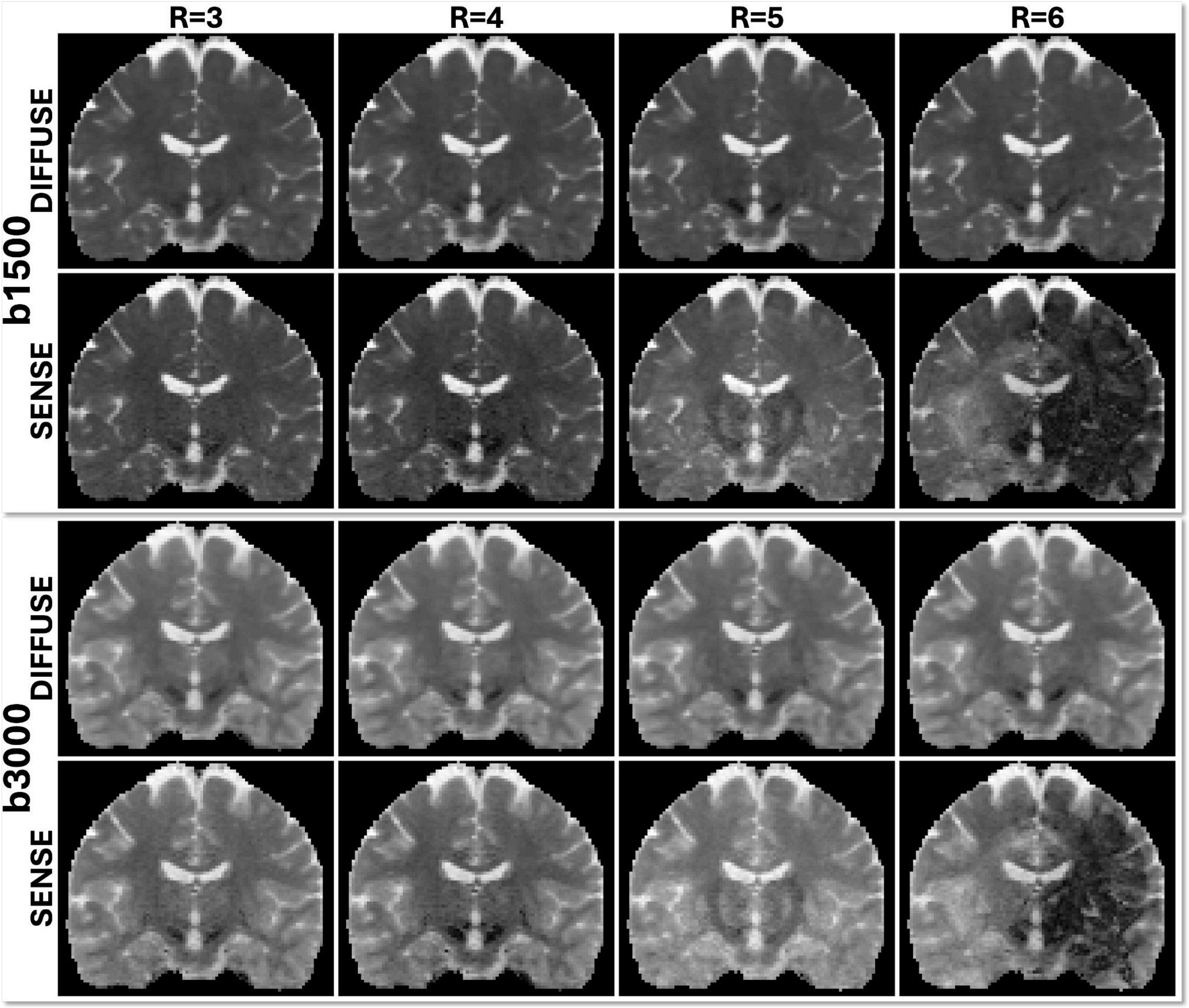


**Fig. S2. Mean diffusivity (MD, scale 0–0.003 mm$^2$/s) for retrospective HCP multi-shell acceleration. Maps are arranged by acceleration factor and shell, following the same layout as the DWI panels and DTI orientation maps.**

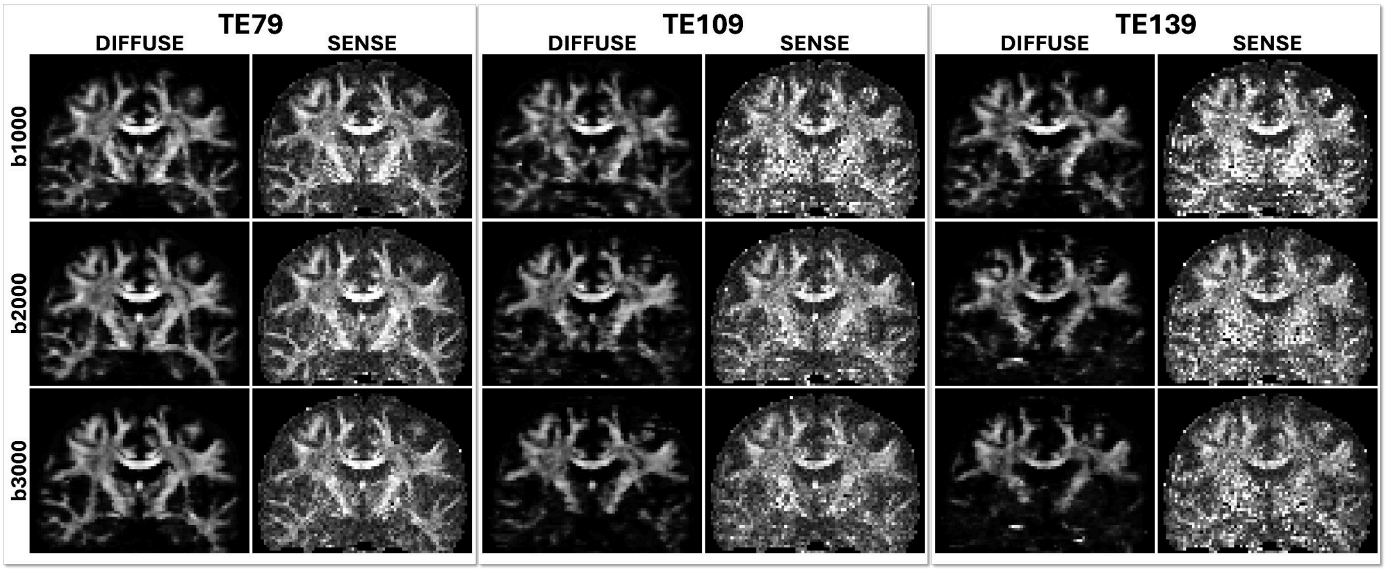


**Fig. S3. Fractional anisotropy (FA, scale 0–0.8) maps at coronal slice $y = 53$ across the complete $3{\times}3$ b-value/TE grid in the multi-shell multi-TE experiment. Rows show $b = 1000$, 2000, and 3000 s/mm$^2$. Column pairs show TE values of 79, 109, and 139 ms, with DIFFUSE and SENSE displayed side by side within each pair.**

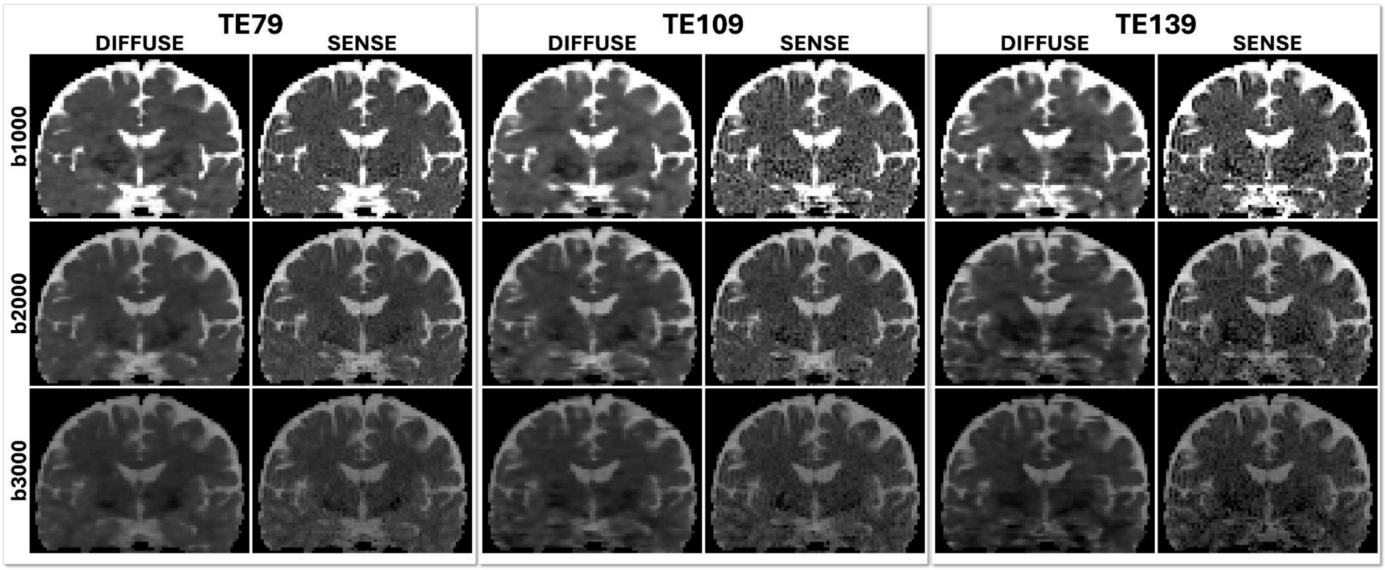


**Fig. S4. Mean diffusivity (MD, scale 0–$2.5 \times 10^{-3}$ mm$^2$/s) maps at coronal slice $y = 53$ across the complete $3 \times 3$ b-value/TE grid in the multi-shell multi-TE experiment. Rows show $b = 1000$, 2000, and 3000 s/mm$^2$. Column pairs show TE values of 79, 109, and 139 ms, with DIFFUSE and SENSE displayed side by side within each pair.**

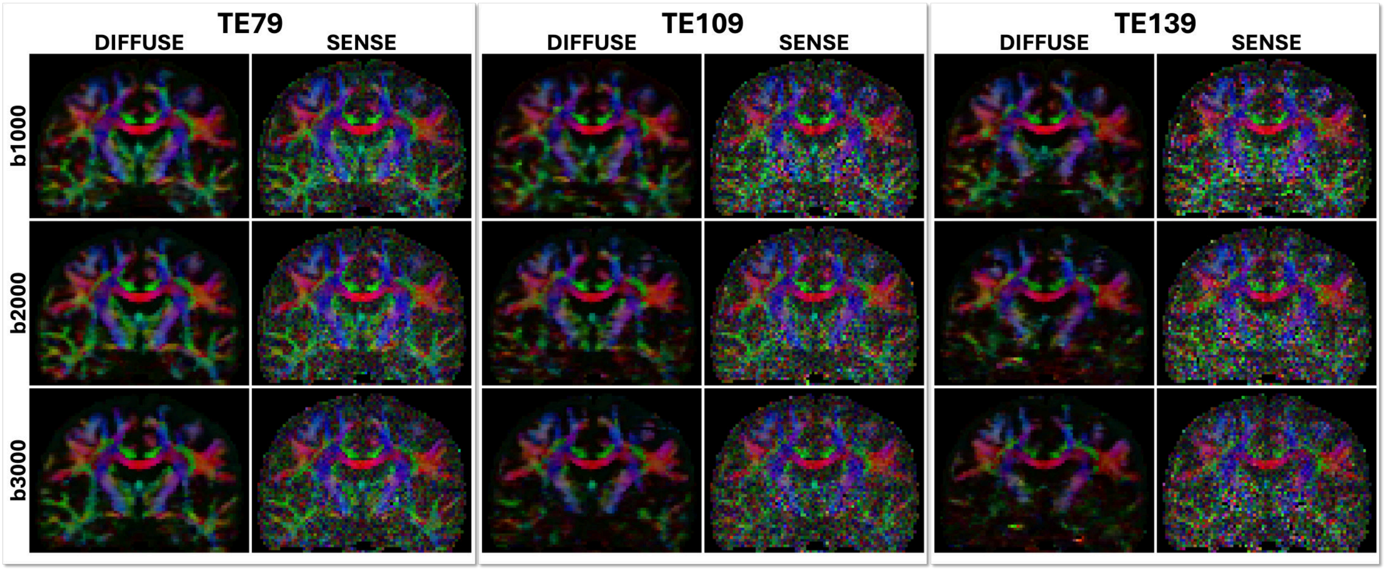


**Fig. S5. DTI orientation maps at coronal slice $y = 53$ across the complete $3\times3$ b-value/TE grid in the multi-shell multi-TE experiment. Rows show $b = 1000$, 2000, and 3000 s/mm$^2$. Column pairs show TE values of 79, 109, and 139 ms, with DIFFUSE and SENSE displayed side by side within each pair.**
